# Cancer-associated fibroblast-derived ROR2 induces WNT/PCP activation and polarized migration in receiving gastric cancer cells

**DOI:** 10.1101/2022.04.07.487474

**Authors:** Sally Rogers, Chengting Zhang, Vasilis Anagnostidis, Melissa Fishel, Fabrice Gielen, Steffen Scholpp

## Abstract

Bone marrow-derived mesenchymal stem cells and cancer-associated fibroblasts in the tumor-stromal environment have been linked to cancer progression in many studies. These fibroblasts provide signaling factors to the tumor cells that promote proliferation, survival, invasion, and metastasis. One signaling pathway influencing tumor cell behavior is the WNT/Planar Cell Polarity (PCP) signaling in gastric cancer. Here, we show that the gastric tumor cell line, AGS, can respond to the PCP ligand WNT5A, however, express a very low level of the bona-fide WNT/PCP receptor, ROR2. At the same time, we find that CAF display long filopodia and had significantly higher levels of ROR2 than normal gastric fibroblasts. By high-resolution imaging, we observe a direct, cytoneme-mediated transfer of a complex containing ROR2 and WNT5A from CAF to the gastric cancer cells. The amount of ROR2 transferred correlated with JNK signaling in receiving cells, showing a direct requirement for receptor transfer. Co-culture of AGS with CAF expressing a dominant-negative form of ROR2 exhibited reduced actin polarization and migration compared to wild-type CAF. Furthermore, induction of migration via paracrine ROR2 transfer was observed in a zebrafish *in vivo* model. These unexpected findings demonstrate a fresh role in the direct transfer of a Wnt receptor from a signal-producing cell to a receiving cell and explain the mechanism by which gastric cancer cells expressing low levels of ROR2 can respond to a WNT5A-high tumor microenvironment.

## Background

Gastric cancer is the fifth most common cancer and the third leading cause of cancer-related deaths worldwide (Jemal et al., 2011). Despite some decline in incidence in recent years, there is still an urgent need to develop novel therapeutic approaches to improve prognosis. The aberrant activation of the Wnt/β-catenin signaling pathway in tumor cells has been implicated in the development and progression of gastric cancer (reviewed in (Chiurillo, 2015) and presents an attractive potential pharmaceutical target. However, the role of the Planar Cell Polarity (PCP) Wnt signaling pathway in gastric tumor cell migration is less well-characterized. Wnt ligands such as WNT5A are considered to activate WNT/PCP upon binding to receptor tyrosine kinase-like orphan receptor 2 (ROR2) or RYK in concert with Frizzled receptors (FZD) (Stricker et al., 2017, Rogers and Scholpp, 2022). Subsequently, WNT/PCP leads to the activation of c-Jun N-terminal kinase (JNK) signaling, one of the best-characterized effectors associated with cell polarity, cytoskeleton rearrangements, and cell migration (Oishi et al., 2003, Minami et al., 2010).

WNT5A expression is associated with several cancers, resulting in constitutive activation of ROR2-mediated signaling and contributing to tumor progression (Enomoto et al., 2009, O’Connell et al., 2010, Ren et al., 2011). In gastric cancer, WNT5A is highly expressed in the stroma (Saitoh et al., 2002), contributes to the tumor microenvironment, and is explicitly upregulated in cancer-associated fibroblasts compared to fibroblasts isolated from the normal gastric stroma (Wang et al., 2016). We have previously shown that Wnt ligands can be transported intercellularly over long distances via specialized actin-rich signaling filopodia, also known as cytonemes. For example, cytoneme-mediated transport of Wnt8a from a signal-producing cell to a signal-receiving cell regulates the patterning of the neural plate in zebrafish gastrula (Stanganello et al., 2015, Mattes et al., 2018; Brunt et al., 2021). More recently, we have shown that Wnt3 is transported intercellularly via cytonemes and facilitates tumor progression in gastric cancer (Routledge et al., 2022). Therefore, cytoneme-mediated intercellular transport of WNT5A from gastric CAF to gastric cancer cells provides an attractive hypothesis to describe how long-range signaling may be achieved in the tumor microenvironment.

However, the identity of the receptor for WNT5A on gastric cancer cells remained unclear, given that ROR2 is commonly down-regulated in these tumors (Astudillo, 2021, Yan et al., 2016). Here, we show that in addition to WNT5A, ROR2 is also upregulated on both pancreatic and gastric cancer-associated fibroblasts compared to the gastric cancer cell line, AGS, and normal gastric fibroblasts. Furthermore, endogenous ROR2 and WNT5A co-localize on cytoneme tips extended by CAF. Surprisingly, we found that WNT5A/ROR2 is transported from CAF to AGS cells as a complex. These receptor/ligand complexes remained stably associated in receiving cells. We then use a reporter system to demonstrate that activation of the JNK signaling pathway in ROR2 receiving AGS cells is proportional to the amount of delivered ROR2. Thereupon, CAF induce actin polarization and directional migration in AGS cells in a ROR2-dependent manner in both *in vitro* and *in vivo* 3D models. Taken together, our results provide evidence of a novel intercellular signaling mechanism, in which both a Wnt ligand and the cognate co-receptor ROR2 are transported from a producing cell to a receiving cell together and activate the WNT/PCP pathway. This observed transport of the receptor explains how gastric cancer cells, low in ROR2, can respond to a WNT5A-high microenvironment.

## Methods

### Cell Culture

The human gastric adenocarcinoma cell line AGS was purchased from ECACC and maintained in Roswell Park Memorial Institute (RPMI) 1640 Medium supplemented with 10% fetal bovine serum (Fisher). Fibroblast cell lines were generously gifted by Dr. Melissa Fishel of IU Simon Cancer Research Center and isolated as described in (Richards et al., 2017). Fibroblast nomenclature was reduced for purposes of simplicity with “pCAF2” referring to UH1303-02 cells. pCAF2 cells were maintained in Dulbecco’s modified eagle medium supplemented with 10% fetal bovine serum. Human primary gastric myofibroblasts were generously gifted by Andrea Varro and are as described in (Holmberg et al., 2012). Gastric myofibroblasts were cultured in Dulbecco’s modified Eagle medium (DMEM) with L-glutamine containing 10% fetal bovine serum, 1% modified Eagle medium nonessential amino acid solution, 1% penicillin/streptomycin, and 2% antibiotic-antimycotic. The medium was replaced routinely every 48 – 60 hours, and cells were passaged at confluence up to 10 times. The stable ROR2 knockout JNK reporter AGS cell line was generated by transfection first with the validated ROR2 CRISPR plasmids sc-401324 (Santa Cruz Biotechnology). ROR2 knockout clones were confirmed by sequencing, and the B18 clone was transfected with the JNK KTR-mCherry plasmid (Regot et al., 2014, Miura et al., 2018), followed by selection with blasticidin. A stable clone was selected, which expressed the JNK reporter at a level suitable for detection by confocal microscopy. All cells were maintained at 37 °C with 5% CO2.

### Plasmids and transfection

The following plasmids were used in transfections: pCS2+ GPI-anchored mCherry (Membrane-mCh) (Scholpp et al., 2009), pCS2+ Gap43-GFP (membrane-GFP), pCS2+ Ror2^3i^ (Mattes et al., 2018), JNK KTR-mCherry (Regot et al., 2014), pCS2+ Ror2-mCherry (cloned in with ClaI and XbaI), pCS2+ Ror2-eBFP2 (cloned by inserting eBFP2 into pCS2+ Ror2 plasmid using XbaI and SnaBI), pCDNA Ror2-eBFP2 (Ror2 and eBFP2 amplified and cloned into pcDNA3.1 using the PmeI and NotI restriction sites), pcDNA dCDRor2-eBFP2 (Ror2 minus intracellular domain amplified and cloned into the pCDNA-eBFP2 backbone using NotI and XbaI restriction sites), pCS2+ Wnt5A-eGFP (Wnt5a cloned into a pCS2+ GFP backbone using ClaaI and XbaI restriction sites). Transfections were performed using a 4D-Nucleofector Unit (Lonza), with the P2 Primary Cell Kit for pCAF2 cells and the SF Cell Line kit for AGS cells. AGS cells were additionally transfected with Fugene HD Transfection Reagent (Promega) if high transfection efficiency was not critical.

### qPCR

RNA for qPCR was collected from cell pellets using QIAGEN RNeasy kit according to the manufacturer’s instructions. RT-qPCR was then performed using SensiFAST™ SYBR® Lo-ROX One-Step Kit with half volumes according to manufacturer’s protocol and run using Applied Biosystems QuantStudio6 Flex. Primer sequences used were as follows: *ROR2* (Forward 5’-GTACGCATGGAACTGTGTGACG -3’); (Reverse 5’-AAAGGCAAGCGATGACCAGTGG -3’), *WNT5A* (Forward 5’-TACGAGAGTGCTCGCATCCTCA -3’); (Reverse 5’-TGTCTTCAGGCTACATGAGCCG-3’),*GAPDH* (Forward 5’-GTCTCCTCTGACTTCAACAGCG -3’); (Reverse 5’-ACCACCCTGTTGCTGTAGCCAA -3’). Relative mRNA expression was calculated using the 2^-ΔΔCt^ method.

### Flow cytometry

Cells were harvested, washed once in PBS, and fixed in 4% PFA in PBS on ice for 15 minutes. Cells were washed in PBS then permeabilized using 0.5% BSA PBS buffer containing 0.3% Triton-X 100 for 10 minutes at room temperature. Cells were washed again, incubated with anti-Wnt5a primary mouse monoclonal antibody clone A-5 (Santa Cruz) for 30 minutes on ice, washed in 0.5% BSA PBS buffer and incubated with goat anti-mouse IgG AF647 (Abcam) for 30 minutes on ice. Cells were washed again and analysed on an Attune NxT flow cytometer (Thermofisher). Data were analysed and presented using FlowJo.

### Western blotting

Cell lysates were collected from cell pellets by resuspending in 50µl of ice cold RIPA lysis buffer (Fisher) containing 1x Halt Protease Inhibitor cocktail (Fisher) per 1×10^6^ cells. Cells were incubated on ice for 30 mins before centrifuging at 12,000RPM, 4°C for 10 minutes and removing the supernatant. A BCA protein assay kit (Pierce) was then used according to manufacturer’s microplate protocol to calculate protein concentrations. For SDS-PAGE gel electrophoresis, required volume of 4x Laemmli buffer (with 10% βME) was added to protein samples and incubated at 95°C for 5 mins. 30µg of protein was then loaded into two BIORAD Mini-PROTEAN pre-cast gels and run at 100V for 90 minutes. Gels were then Western-blotted to PVDF membrane, and blocked in 5% bovine serum albumin for 2 hours at RT. Primary antibodies (ROR2: Cell Signalling Technologies D3B6F, GAPDH: Merck 6C5) were added and incubated on a roller overnight at 4°C. The following day, membranes were washed 3x with TBS-T before incubating in secondary antibodies goat anti-mouse IgG H&L (GAPDH) and goat anti-rabbit IgG H&L (ROR2) (Abcam, both IRDye® 800CW) in TBS-T for 1hr at RT. Membranes were then washed 3x with TBS-T and 3x with TBS before imaging on LiCor Odyssey CLx and processing on ImageStudio software.

### Immunofluorescence

Cells were plated onto glass coverslips and following 24 hours incubation, were washed in 1xPBS and fixed using modified MEM-Fix (4% formaldehyde, 0.25-0.5% glutaraldehyde, 0.1M Sorenson’s phosphate buffer, pH7.4) (Rogers and Scholpp, 2021, Bodeen et al., 2017) for 7 minutess at 4°C. Cells were then incubated in permeabilization solution (0.1% TritonX-100, 5% serum, 0.3M glycine in 1xPBS) for 1hr at RT. Primary antibodies (mouse anti-Ror2: Santa Cruz H-1, rabbit anti-Wnt5a/b: ProteinTech 55184-1-AP) were diluted in incubation buffer (0.1% Tween20, 5% serum in 1xPBS) and coverslips incubated in 50 µl spots on parafilm overnight at 4°C. Coverslips were then washed with 1xPBS 3x for 5 minutes before incubation in 50 µl spots of secondary antibodies (goat anti-mouse AF647: Abcam ab150115, goat anti-rabbit AF405: abcam 175652) diluted in incubation buffer for 1hr at RT. Coverslips were then washed 3x for 5 minutes with 1xPBS before mounting onto glass slides using ProLong Diamond anti-fade mountant (Invitrogen) and left to dry for 24hrs before imaging. Confocal microscopy for immunofluorescent antibody imaging was performed on a Leica TCS SP8 laser-scanning microscope using the 63x water objectives.

### JNK reporter assay

AGS *ROR2* KO cells stably expressing JNK-KTR-mCherry (AGS-B18) were co-cultured with either pCAF2 cells or AGS cells transfected with indicated plasmids for 24 hours. Cells were imaged on a Leica TCS SP8 confocal microscope using a 40x oil objective. Fluorescence intensity of each receiving cell’s cytoplasm and nucleus was measured using Fiji software, and cytoplasmic:nuclear ratio (C:N) was calculated.

### Actin Polarization

AGS cells were transfected with LifeAct-GFP and incubated for 24 hours for recovery. pCAF2 cells were transfected as required and also incubated for 24 hours recovery. Cells were trypsinised, counted and the two populations co-cultured for 24 hours. Cells were imaged using a Leica TCS SP8 laser-scanning microscope using the 63x oil objective. The fluorescence intensity was analysed in AGS cells that were in contact with a pCAF2 cell using FIJI. A line was drawn through the middle of the nucleus of the receiving AGS cell to define facing and opposing sides, and the mean fluorescence at the membrane of each side was measured.

### Microfluidic chip fabrication

Microfluidic device fabrication was done using standard soft lithography techniques. Negative photoresist SU-8 3035 (MicroChem, Newton, MA) was spin-coated onto a silicon wafer at 4,500 rpm to a thickness of 30 µm. Pattern cross-linkage occurred via exposure to UV light through chrome mask (Plate size x Thickness: 4”x4” x 0.09”, ARC Quartz) (Micro Lithography Services Ltd). The microfluidic designs were drawn using AutoCAD 2021 (Autodesk). The device consists of four inlets for media/cell induction and a large chamber that is separated in half by a row of a diamond-shaped grid of pillars (40 µm x 40 µm) (Supplementary Figure 3). The pillars are spaced out with a 4 µm gap. Uncured polydimethylsiloxane (PDMS) (Sylgard 184) was poured onto the master (10:1 polymer to cross-linker mixture), degassed, and baked at 70 °C for 4 hours. The PDMS mould was then cut and peeled from the master. Four holes were punched using a 6 mm biopsy punch (Stiefel). The PDMS mould was then plasma bonded (Diener Zepto) to a thin cover slip (22 × 50 mm, 0.13 – 0.17 mm thick). Sterile PBS was immediately flushed through the device to increase the hydrophilic properties of PDMS. The device was placed under UV for 30 minutes prior to loading of samples.

### Microfluidic chip assay

A bespoke microfluidic device was designed to co-culture two cell types in two separate chambers divided by diamond-shaped grid with small 4 µm gaps preventing cells from migrating across while allowing filopodia to establish cell-to-cell communication (supplementary Figure 2). In short, pCAF2 cells were transfected as required and cultured for 24 hours prior to counting, resuspended in DMEM + 5% FBS minus phenol red, and loaded on to one side of the microfluidic chambers. Following a further 24 hours of culture in which the pCAF2 adhered to the glass coverslip, the media was removed from the chamber, and AGS cells transfected with LifeAct-GFP were loaded onto the other side of the chamber in DMEM + 5% FBS minus phenol red. Cells were co-cultured in the chip for 18-24 hours, then live imaged using a Leica SP8 confocal microscope with a 100x oil objective.

### Wound healing assay

AGS or pCAF2 cells were transfected as required and incubated for 24 hours. Cells were then trypsinised, counted and plated into silicone 2 well culture inserts (Ibidi). Following a further 24 hours incubation, cells were treated with 10µg/ml mitomycin C (MMC) for 90 minutes, washed well, and then the culture insert removed. Cells were incubated with live imaging on a Leica DMi8 light microscope with an Ibidi environmental chamber controlling temperature and carbon dioxide. Area of the wound at 18 hours was analyzed using FIJI.

### 3D Invasion assay

pCAF2 cells were transfected as required and cultured for 24 hours for recovery. AGS cells were transfected with pCS2+ Gap43-GFP (membrane-GFP) and also incubated for 24 hours. The 3D invasion assays were performed using GrowDex® hydrogel (UPM, Helsinki, Finland) as the matrix supporting 3-dimensional cell growth. For the experiment, 1×10^5^ pCAF2 cells were resuspended in 40µl complete cell culture medium (DMEM + 5% FBS, minus phenol red), then mixed with 90µl of the 1.0% hydrogel stock to achieve a 0.75% w/v hydrogel solution. 80µl of this suspension was dispensed to a well of a 96 well-transwell plate (Corning) using a wide-bore pipette. The insert was placed on top, and 5×10^3^ transfected AGS cells aliquoted in 100µl of DMEM + 5% FBS minus phenol red on top of the 8 µm pore-size membrane. Media was exchanged once per day for 72 hours, and on the third day, cells were imaged using a Leica DMI6000 light microscope with a 20x dry objective. 1mm stacks were obtained, with a frame at every 5 µm. Images were analysed using Imaris, with green AGS cells converted to spots and the depth of their position in the hydrogel measured. Three stacks were obtained in each well, and the experiments were performed in triplicate.

### In vivo zebrafish migration assay

WIK wild-type zebrafish (*Danio rerio*) were maintained at 28°C and on a 14hr light/10hr dark cycle (Brand et al., 2002). Zebrafish care and all experimental procedures were carried out in accordance with the European Communities Council Directive (2010/63/EU) and Animals Scientific Procedures Act (ASPA) 1986. In detail, adult zebrafish for breeding were kept and handled according to the ASPA animal care regulations and all embryo experiments were performed before 120 hrs post fertilization. Zebrafish experimental procedures were carried out under personal and project licenses granted by the UK Home Office under ASPA, and ethically approved by the Animal Welfare and Ethical Review Body at the University of Exeter. Embryos were injected with membrane mCherry mRNA +/- Ror2 mRNA into 1 cell out of 8 blastomeres, and the adjacent cell was injected with Gap43-GFP mRNA to generate two independent clones in the same embryo. The live embryos were mounted and imaged at 6 hours post-fertilization using a Leica TCS SP8 confocal microscope, using 10x dry objective. The area of the embryo containing red and green cells was determined using FIJI. The ordinary one-way ANOVA together with Tukey’s multiple comparisons test was performed using GraphPad Prism 9.0.

## Results

### Endogenous ROR2/WNT5 complexes are expressed on fibroblast cytoneme tips

WNT/PCP signaling plays a pivotal role in tumor progression in many cancers. However, it is unclear which cells express the PCP ligands and their receptors in the microenvironment. Therefore, we analyzed the expression of one of the most important PCP ligands, WNT5A. We find that WNT5A expression is higher in gastric cancer-associated fibroblasts than in gastric tumor cells and surrounding normal fibroblasts, which has been suggested previously (Wang et al., 2016). *WNT5A* mRNA was shown to be 357-fold higher in primary gastric Normal Myofibroblasts (gNM) compared to AGS cells and over 800-fold higher in primary gastric cancer-associated myofibroblasts (gCAM) and pancreatic cancer-associated fibroblasts (pCAF2) compared to AGS cells using qRT-PCR (Figure 1a). A flow cytometry-based analysis validated this finding at the protein level (Figure 1b). However, previous reports indicate that the cognate WNT5A receptor ROR2 is downregulated in gastric cancer cells (Astudillo, 2021, Yan et al., 2016). Therefore, the primary aim of this study was to further investigate the distribution of WNT5A receptors in a gastric tumor to understand how gastric cancer cells respond to a WNT5A-high environment. To this end, *ROR2* gene expression was quantified using qRT-PCR. We found that *ROR2* expression is significantly increased in gCAM and pCAF2 compared to both AGS and gNM (Figure 1a). Furthermore, the expression of *ROR2* mRNA varied between gCAM isolated from different patients but was higher overall in cancer-associated fibroblasts than gNM, regardless of origin. We confirmed that mRNA expression correlated with protein expression using Western blotting (Supplementary Figure 1). Our previous studies have identified several WNT signaling components localized to signaling filopodia, also known as cytoneme (Stanganello et al., 2015; Mattes et al., 2018; Brunt et al., 2021). We, therefore, established an IF method that fixes these filopodia to examine the cellular localization of endogenous ROR2 and WNT5A (Rogers and Scholpp, 2021). Co-localizing complexes of ROR2 and WNT5A were observed along the length of pCAF2 filopodia, including at the tips of these structures (Figure 1c).

**Figure 1.**
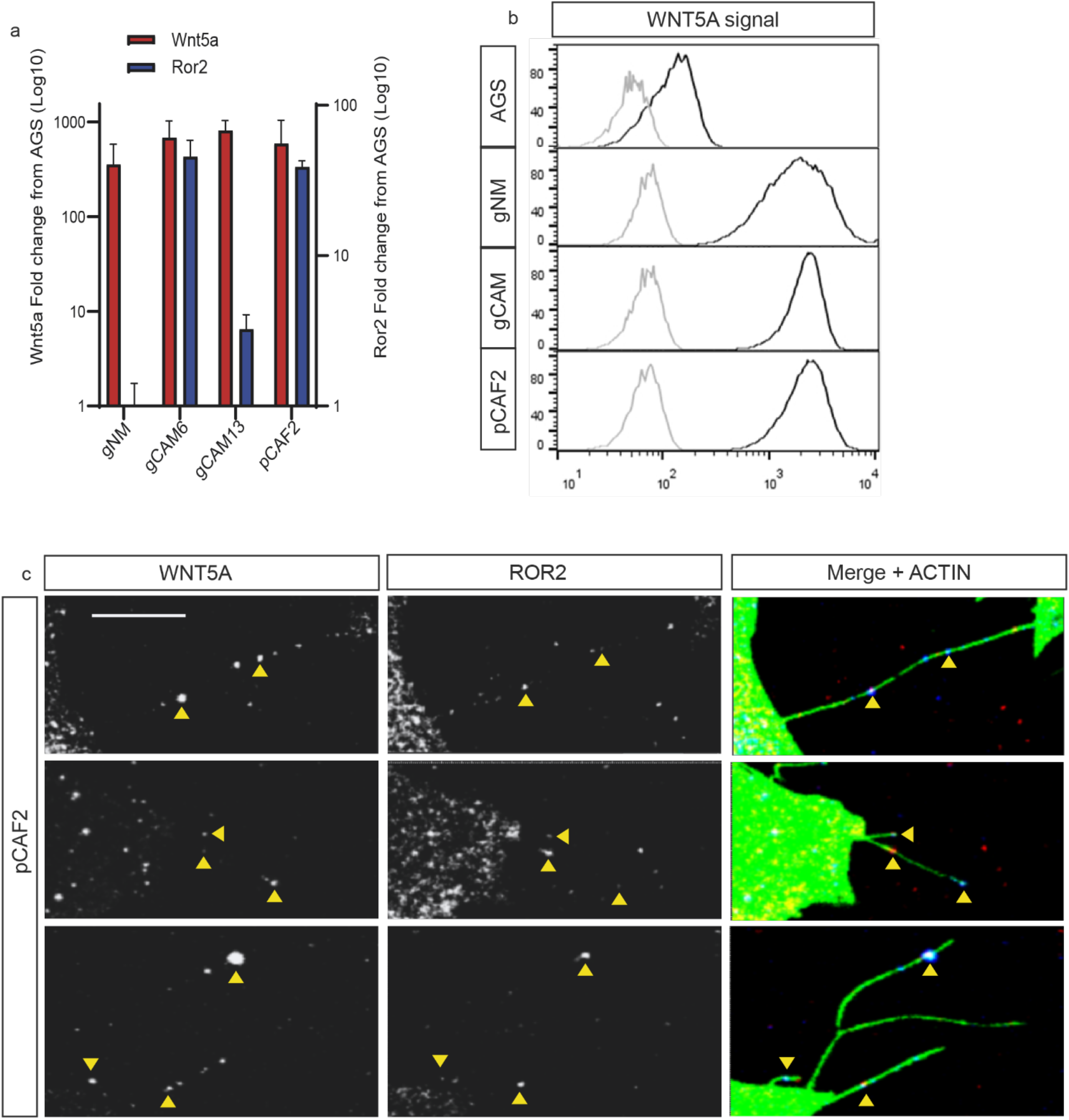
Expression and localization of WNT5A/ROR2 complexes in GC cells. **a**. mRNA levels of the indicated gene shown as fold change away from AGS. Wnt5a is shown in red, values on the left y-axis. Ror2 is shown in blue, values on the right y-axis. gNM - primary gastric Normal Myofibroblasts, gCAM6 – primary gastric cancer-associated myofibroblast patient 6, gCAM13 - primary gastric cancer-associated myofibroblast patient 13, pCAF2 – pancreatic cancer-associated fibroblast cell line 2. Values shown were calculated using the 2^-ΔΔCt^ method and are plotted on a log10 scale. **b**. Protein expression of WNT5A as detected using flow cytometry. Cells as indicated on left panels. The grey histogram indicates control staining, and the black histogram indicates WNT5A staining. **c**. Immunofluorescence on pCAF2 cells fixed using a previously described method to preserve filopodia. The left-hand column indicates WNT5A/B antibody staining, the center panel indicates ROR2 antibody staining, and the right-hand panel indicates WNT5A/B and ROR2 antibody staining (blue and red, respectively) merged with Phalloidin FITC shown in green. Three representative images are shown. The scale bar represents 10 µm.

### ROR2/WNT5 complexes are transferred from CAF to AGS cells via cytonemes

Our initial hypothesis was that WNT5A produced by CAF (signal-producing cells) would bind to the low levels of ROR2 on AGS cells (signal-receiving cells). However, to our surprise, when fluorescently labeled ROR2 and WNT5A were over-expressed together in AGS cells, receptor-ligand complexes were observed not only co-localizing at cytoneme tips but also co-localizing in neighboring, untransfected cells (Supplementary Figure 2a). Moreover, when a fluorescent membrane marker was expressed alongside ROR2 and WNT5A in transfected AGS cells, all three components were observed co-localizing in receiving AGS cells (Figure 2a). We observed the same phenomena occurring between pCAF2 cells over-expressing WNT5A and ROR2 (Supplementary Figure 2b), suggesting that cytoneme-based delivery is a common mechanism within a tumor tissue. Time-lapse images revealed that WNT5A/ROR2 complexes can be directly transferred via both AGS and pCAF2 producing cell cytonemes to a receiving AGS cell (Figures 2b and c, respectively). Furthermore, these receptor-ligand complexes remain stable and continue to co-localize in AGS receiving cells for a considerable amount of time. Taken together, these results suggest that the tip of the producing cell cytoneme buds off and is endocytosed by the receiving cell. We hypothesized that in this way, the ROR2-low cells, like the gastric tumor cells AGS, could increase their responsiveness to fibroblast-produced WNT5A as they are receive a functional receptor.

**Figure 2.**
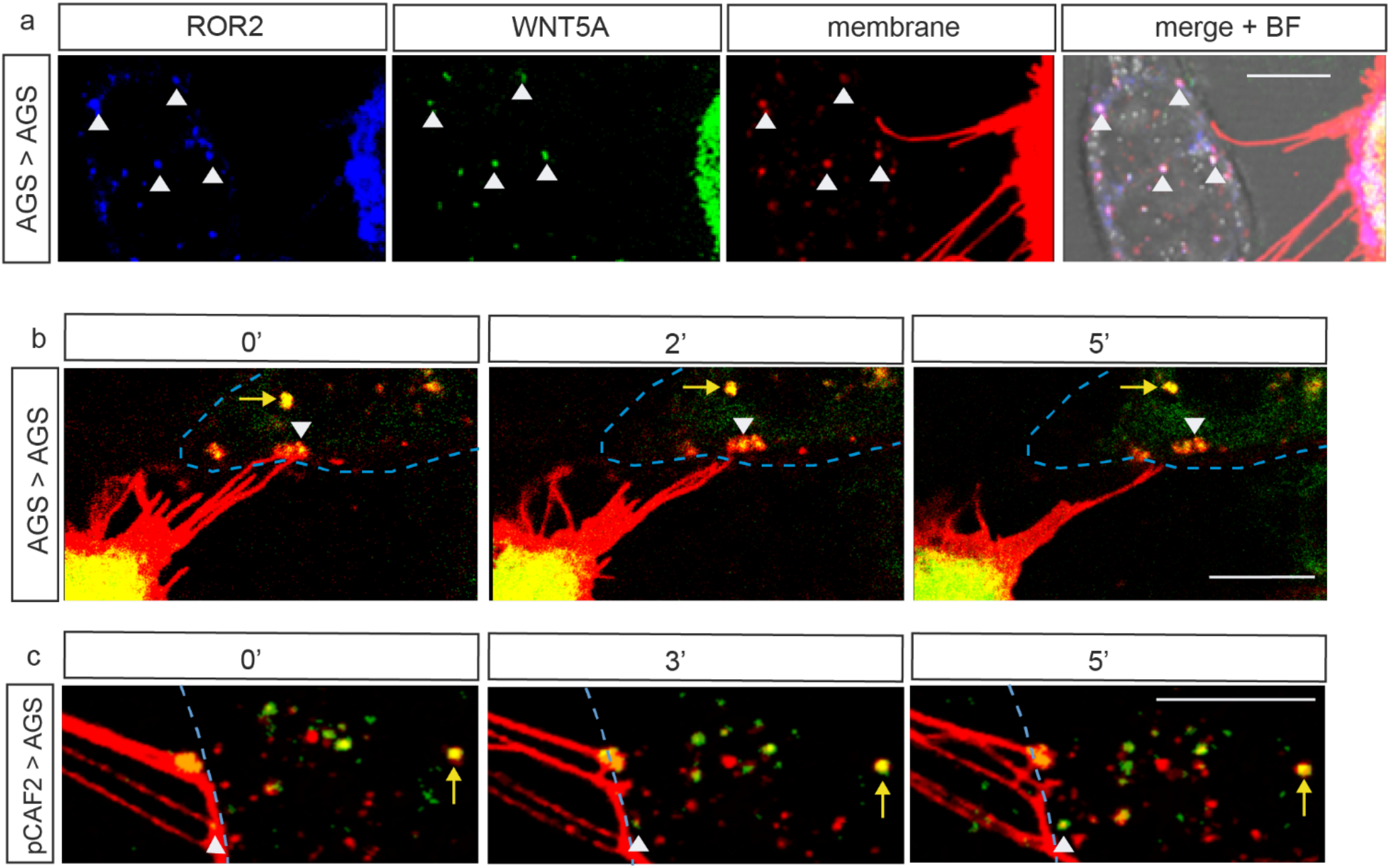
Transfer of ROR2 via producing cell cytonemes to receiving cells in gastric cancer. **a**. Confocal image of an AGS cell transiently transfected with ROR2-BFP, WNT5A-GFP, and membrane mCherry. ROR2/WNT5A/membrane complexes in a receiving, untransfected AGS cell are indicated by white arrows. The scale bar represents 10 µm. **b**. AGS cell expresses ROR2-mCherry and WNT5A-GFP on cytoneme tips (white arrow), which can be seen contacting a neighboring AGS receiving cell indicated by the blue dashed line (see also supplementary figure 1a). Time-lapse images, as indicated, show a ROR2/WNT5A complex leaving the cytoneme tip and co-localizing in the receiving cell. Other ROR2/WNT5A complexes from the producing cell can localize in the receiving cell, as indicated by the yellow arrow. **c**. pCAF2 cell expresses ROR2-mCherry and WNT5A-GFP on cytoneme tips (white arrow), which can be seen contacting a neighboring receiving AGS cell indicated by the blue dashed line (see also supplementary figure 2b). Time-lapse images, as indicated, show a ROR2/WNT5A complex leaving the cytoneme tip and co-localizing in the receiving cell. Other ROR2/WNT5A complexes from the producing cell can be seen localizing in the receiving cell, as indicated by the yellow arrow.

### CAF produced ROR2 can induce Jnk signaling in recipient AGS cells

Next, we used a JNK signaling reporter system to quantify the PCP response in receiving cells to test our hypothesis. To determine the response of AGS cells to transferred ROR2 (rather than endogenous receptor), a ROR2 knockout AGS cell line was produced, which was subsequently transfected with the JNK-KTR-mCherry reporter plasmid, and a stable clone was selected. This cell line (AGS-B18) was used as the receiving cells, in which the reporter translocates from the nucleus to the cytoplasm upon activation of the JNK signaling pathway (Figure 3a). The pCAF2 cell line was used as the producing cell in consequent experiments, as they have similar expression levels of ROR2 and WNT5A as the primary gCAM but survive longer and are genetically and phenotypically constant.

**Figure 3.**
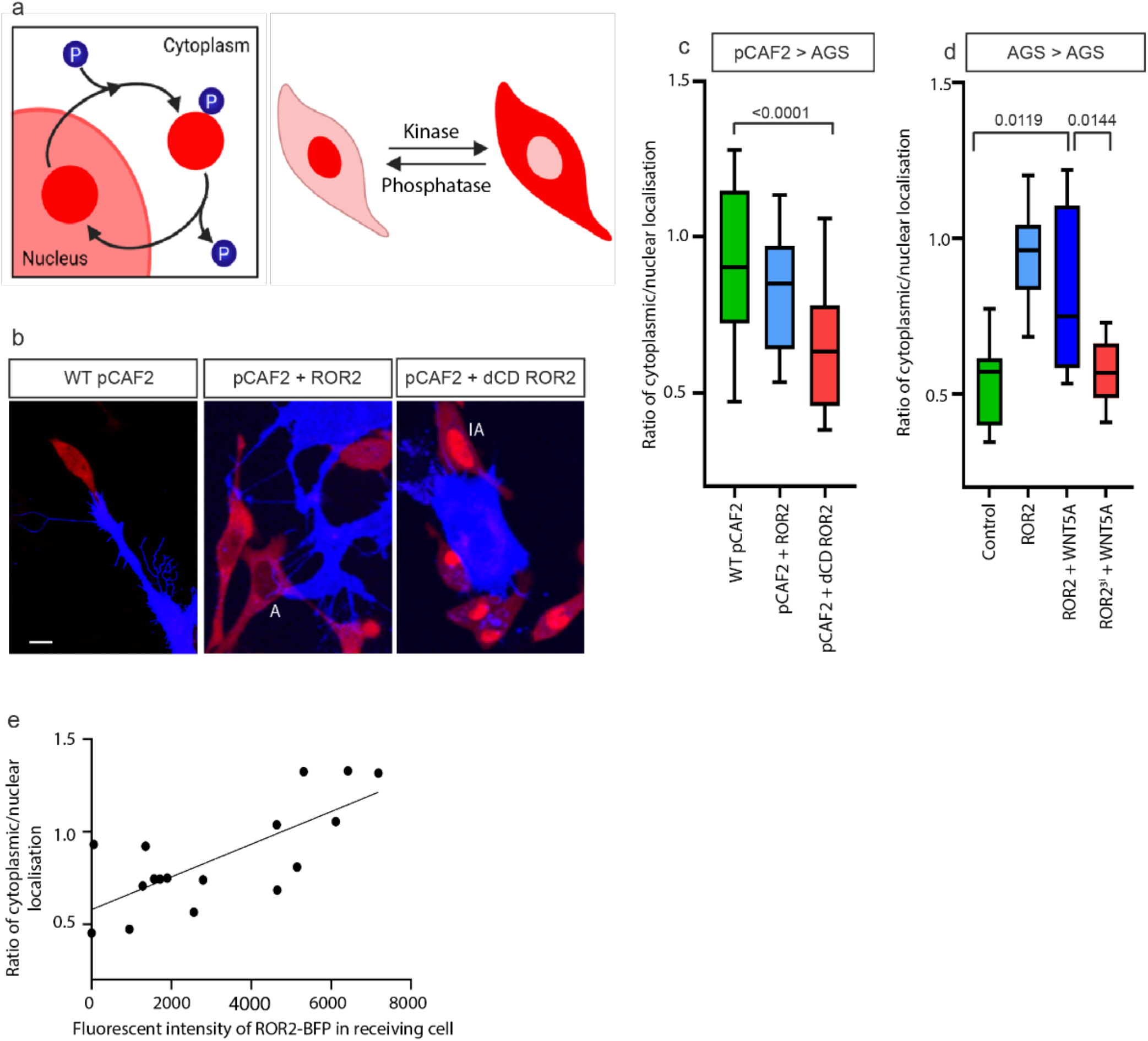
ROR2 induces JNK signaling in receiving AGS cells. **a**. Cartoon depicting JNK signaling assay (adapted from (Regot et al., 2014). Upon activation of the JNK signaling pathway, the mCherry reporter protein translocates from the nucleus to the cytoplasm of the cell. **b**. Representative confocal images of AGS-B18 cells co-cultured with wild-type pCAF2 transfected with either membrane GFP (left panel), ROR2 BFP (center panel), or dCD ROR2 BFP (right panel). The scale bar represents 10μm. **c**. Quantification of the cytoplasmic to nuclear ratio of JNK reporter signal in receiving AGS-B18 cells co-cultured with pCAF2 cells as described in b. Boxes represent 95% quartile, the centerline indicates the mean, and whiskers indicate the range. **d**. Quantification of the cytoplasmic to nuclear ratio of JNK reporter signal in AGS-B18 cells co-cultured with AGS cells transfected with either membrane GFP (control), ROR2, ROR2, and WNT5A, or dominant negative ROR23i plus WNT5A as indicated. **e**. The intensity of ROR2-BFP was quantified in AGS-B18 receiving cells following co-culture with pCAF2 transfected with ROR2-BFP. Values were plotted against corresponding cytoplasmic/nuclear JNK reporter localization, and a simple linear regression was performed.

pCAF2 cells were transfected with either a membrane marker (WT pCAF2), ROR2, or a dominant-negative ROR2 lacking the intracellular tyrosine kinase domain (dCD ROR2). These were co-cultured with AGS-B18 cells for 24 hours, then imaged using confocal microscopy. Receiving cells for downstream analysis were determined as those in direct contact with a transfected pCAF2 filopodium (Figure 3b). Quantification of the cytoplasmic/nuclear ratio of reporter protein indicated that the Jnk signaling pathway is activated in receiving AGS cells that are co-cultured with both WT pCAF2 and pCAF2 overexpressing ROR2. However, JNK signaling is significantly decreased in receiving AGS cells that are co-cultured with pCAF2 cells overexpressing dCD ROR2 (Figure 3c).

To further examine the contribution of producing cell ROR2 to the induction of JNK signaling in the receiving cell, AGS cells were used as the producing cell line. AGS cells are lower in endogenous ROR2 and WNT5A than pCAF2 cells (figure 1). This strategy limits the effect of other factors produced by the fibroblasts. Co-culture of the AGS-B18 cells with ROR2/WNT5A transfected producing AGS cells resulted in a significant increase in paracrine JNK activation. This effect was ablated when the producing AGS cells were transfected with a dominant-negative form of ROR2 containing inactivating mutations in the tyrosine kinase domain (ROR23i) together with WNT5A (Figure 3d). Furthermore, the amount of transferred ROR2-BFP was quantified in a subset of AGS-B18 receiving cells co-cultured with pCAF2 transfected with ROR2 BFP. Receiving cells were included for analysis in this subset when the transferred ROR2 was quantifiable and the JNK reporter activity was within a linear range. Simple linear regression was used to test if the amount of ROR2 transferred significantly predicted the ratio of JNK localization (Figure 3e). The overall regression was statistically significant (R2 = 0.5471, F (df regression, df residual) = 18.12, p = 0.0007), suggesting a direct correlation between transferred ROR2 and JNK signaling activation. This result suggest the active ROR2 receptors are transported on cytonemes to the receiving cells to induce WNT/PCP signalling in a paracrine fashion..

### CAF-produced ROR2 induces polarization of the actin cytoskeleton in recipient AGS cells

Activation of JNK signaling via the WNT/PCP pathway is associated with cell polarity and migration. To determine whether transferred ROR2 can induce a polarized actin distribution in receiving cells, AGS cells were transfected with LifeAct-GFP and co-cultured with pCAF2 cells expressing either ROR2 or dCD ROR2. Individual receiving AGS cells contacted by a pCAF2 cell were imaged (Figure 4a-c). The ratio of fluorescent membrane actin on the facing side of the cell over the opposing side of the contact site was measured (Figure 4d). We found a significant polarization of actin to the side of AGS cells in contact with pCAF2 cells expressing ROR2, but not dCD ROR2, compared to control AGS cells cultured in the absence of pCAF2 cells.

**Figure 4.**
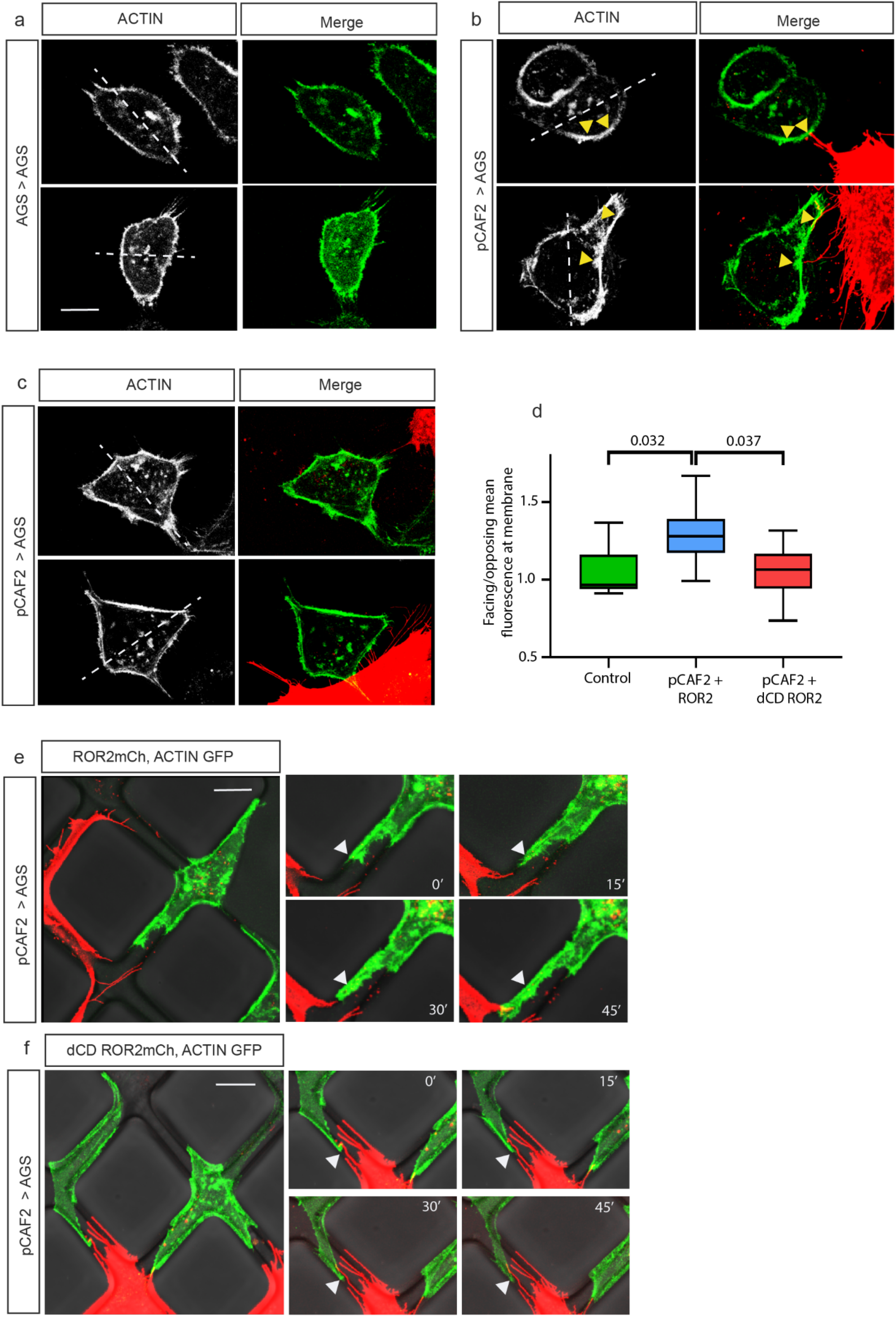
ROR2 induces actin polarization in receiving AGS cells. AGS cells were transfected with LifeAct GFP and cultured either **a**) on their own (Control) or with pCAF2 cells overexpressing **b**) ROR2 mCherry or **c**) dCD ROR2 mCherry. Cells were imaged using confocal microscopy. The left-hand panel shows the 488nm channel only (LifeAct-GFP), and the right-hand panel shows the merge of the green channel with the red channel to show the location of the producing cell and contact points. White dashed lines indicate a bisecting line through the center of the nucleus used to define the facing and opposing sides of the receiving cell. Yellow arrows indicate actin accumulation at contact points with ROR2 producing pCAF2 cells. Two representative images from each condition are shown. **d**. Quantification of facing/opposing mean fluorescent-labeled actin in receiving AGS cells in contact with cells as indicated on the x-axis. **e** and **f**. pCAF2 cells transfected with either ROR2 mCherry or dCD ROR2 mCherry, respectively, were co-cultured with LifeAct-GFP expressing AGS cells in a bespoke microfluidic device enabling cell communication via filopodial networks. Time-lapse images were taken using a confocal microscope, as shown on the right-hand side, at indicated time points in minutes. The white arrow indicates the starting location of the extended AGS cell in each frame. The scale bar represents 10µm.

Due to the complex nature of the pCAF2 filopodial network, we developed a PDMS microchamber to further explore the effect of CAF-derived ROR2 on AGS migration. AGS and pCAF2 cells are cultured in chambers on either side of a series of diamond-shaped pillars, providing better separation and directionality to visualize the molecular interactions between the two populations of cells. Pillars with a dimension of 40μm x 40μm and gaps of 4μm retain the pCAF2 cells while allowing filopodial contacts to form and the AGS cells to migrate in two dimensions (Supplementary Figure 3). AGS cells were transfected with LifeAct-GFP and co-cultured in the PMDS microchambers with pCAF2 cells expressing ROR2 mCherry. After 20 hours of incubation, we observed polarization of actin in individual AGS cells towards the pCAF2 cells confirming our earlier experiments. Within 45 minutes, we observed migration toward the ROR2 producing cell (Figure 4e). No migration was observed when pCAF2 cells expressing dCD ROR2 mCherry were co-cultured with the receiving AGS cells (Figure 4f).

### ROR2 induces directional migration and invasion in 3D models

To determine whether the ROR2-dependent induced migration of individual receiving AGS cells observed in the microchambers applied to whole populations of AGS cells, we utilized a standard 2D migration/wound healing assay. Surprisingly, we found that populations of AGS cells transiently expressing ROR2 and WNT5A had a larger remaining gap after 18 hours of incubation than wild-type AGS cells (Supplementary Figures 4a and 4b). However, upon close examination of time-lapse series of wild-type AGS co-cultured with pCAF2 cells, we observed that AGS cells in close proximity to pCAF2 cells migrate within a narrow radius of the pCAF2 cell rather than migrating away and across the gap (Supplementary Figure 4c). In contrast, AGS cells in the same field that did not start close to a pCAF2 cell migrated much further. This led us to hypothesize that the pCAF2 cells induce a directional, polarized migration in the AGS cells rather than a general increase in migratory capacity. To test this, we established an advanced 3D invasion assay (Figure 5a). In this assay, pCAF2 transfected with either ROR2 or dCD ROR2 were cultured in Growdex hydrogel in 96-well plates, and AGS cells transfected with membrane GFP were cultured in a monolayer in transwells placed on top of, and in contact with, the hydrogel. Following incubation for 72 hours, the wells were imaged using a fluorescent microscope, and the depth that individual AGS cells migrated into the hydrogels was measured using Imaris (Figure 5b). AGS cells migrated significantly less distance into the hydrogel containing pCAF2 cells expressing dCD ROR2 than either wild-type pCAF2 cells or pCAF2 cells overexpressing ROR2 (Figure 5c). This adds further evidence to support our hypothesis that the pCAF2 cells provide a ROR2-dependent signal to the AGS cells to invade and migrate in a directional fashion.

**Figure 5.**
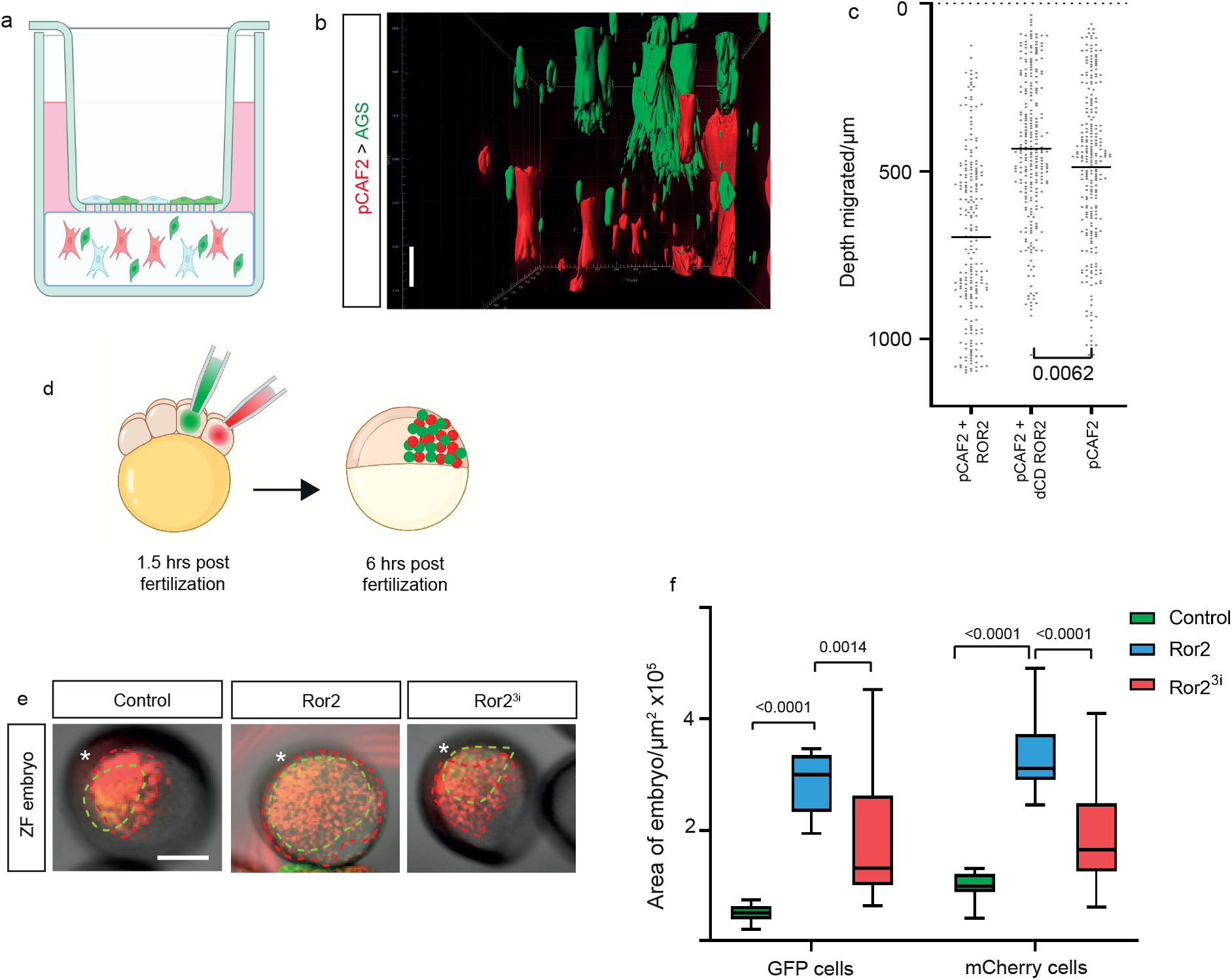
ROR2 induces polarized migration and invasion in 3D models. **a**. Cartoon demonstrating the 3D invasion assay. AGS cells transiently transfected with membrane GFP are cultured in a monolayer in transwells with an 8μm pore size. pCAF2 cells transfected with either membrane mCherry, ROR2 mCherry, or dCD ROR2 mCherry are cultured in 3D using Growdex at the bottom of the well ensuring contact is made between the Growdex and the transwell. **b**. Following incubation for 72 hours, the Growdex is imaged using fluorescent light microscopy, and images are analyzed in Imaris. The scale bar represents 50µm. **c**. depth into the hydrogel that GFP+ive AGS cells invade when cultivated with pCAF2 cells transfected with either membrane mCherry (control), ROR2 mCherry or dCD ROR mCherry as indicated on the x-axis. Points represent individual AGS cells and include technical and biological repeats. The dashed line at the top of the graph represents the location of the transwell. Bars represent the geometric mean for each condition. **d**. Embryos were injected with membrane mCherry mRNA only (control) or plus Ror2 mRNA (Ror2) or Ror2^3i^ mRNA (Ror2^3i^) into 1 cell out of 8 blastomeres, and the adjacent cell was injected with Gap43-GFP mRNA to generate two independent clones in the same embryo. **e**. The live embryos were mounted and imaged at 6 hours post-fertilization, and the area of each embryo containing red and green-labeled cells was measured for each condition as a representation of how far cells from each clone (*) had migrated. Scale bar represents 200µm. **f**. Area of embryos containing GFP or mCherry positive cells indicated on the x-axis for the different conditions (green bars: control – injected with membrane mCherry alone, blue bars: injected with Ror2 mRNA and membrane mCherry, red bars: injected with Ror2^3i^ and membrane mCherry). Boxes represent 95% quartile, the centerline indicates the mean, and whiskers indicate the range. Significance was calculated using ordinary one-way Anova with Tukey’s multiple comparison test.

Next, we performed an *in vivo* migration experiment in zebrafish to further support our idea that Ror2 expressing cells increase directional migration of the neighboring cells. Therefore, wild-type zebrafish embryos were injected with membrane mCherry mRNA +/-Ror2 mRNA into 1 cell out of 8 blastomeres, and the adjacent cell was injected with Gap43-GFP mRNA. Therefore, we generated two independent clones in the same embryo (Figure 5d). The live embryos were mounted and imaged at 6 hours post-fertilization (hpf) (Figure 5e). The area of the embryo containing red and green cells was measured. mCherry expressing cells migrated over a greater area of the embryo when co-expressing ROR2, suggesting that autocrine PCP signaling drives this migratory process (Figure 5f). This effect was ablated when the mCherry cells co-expressed dominant negative ROR23i. Interestingly, the GFP expressing cells also migrated over a greater area of the embryo when mCherry plus ROR2 was expressed in the adjacent cell population compared to embryos containing clones of mCherry alone cells or mCherry plus ROR23i (Figure 5e). This indicates that the GFP expressing cells are induced to migrate further, presumably via paracrine transfer of ROR2 from adjacent cells.

Collectively, these results indicate that the transfer of the ROR2 receptor from cancer-associated fibroblasts to gastric cancer cells increases the capacity of the receiving tumor cell to respond to WNT/PCP signaling - even in the absence of endogenous ROR2 receptors. Furthermore, transferred and active Ror2 receptors affect polarity and directional migration in the cancer cell and increase migration and invasion in 3D *in vitro* and *in vivo* models.

## Discussion

The overexpression of WNT5A in intestinal tumor microenvironments is well documented. Still, the question of how cancer cells respond to PCP signaling is perplexing, given that down-regulation of the concomitant co-receptor ROR2 is observed in many gastric tumors. Here, we show that the WNT5A co-receptor ROR2 can be transferred from cancer-associated fibroblasts to AGS cells, thereby inducing JNK signaling, actin polarization, and directional migration in the receiving cell. This unexpected intercellular signaling mechanism of receptor-ligand complex transfer provides insights and a possible explanation of how gastric cancer cells respond to high WNT5A in their environment.

Frequent upregulation of *WNT5A* mRNA has been previously described in primary gastric cancer (Saitoh et al., 2002), and WNT5A has been shown to be the only Wnt ligand that is consistently upregulated in gCAM (Wang et al., 2016). We confirmed the overexpression of WNT5A in gCAM. In addition, we report the novel finding that WNT5A is equally highly expressed in pancreatic CAF. As with gastric cancer, pancreatic cancer has a poor prognosis, therefore, any insights into the molecular mechanisms of this disease are also valuable. In addition, we identified the upregulation of ROR2 in cancer-associated fibroblasts from both the pancreas and stomach carcinomas compared to normal gastric primary fibroblasts and AGS cells. The gCAM used in our study has been reasonably well-characterized (Holmberg et al., 2012), and upregulated ROR2 has not been previously identified in these primary cells. However, the previous study compared the proteome of gCAM to adjacent tissue-matched (ATM) myofibroblasts, in contrast to the normal myofibroblasts comparison reported here, which may explain why ROR2 was not identified as differentially expressed. For example, the adjacent tissue of patient 6 was defined as intestinal metaplasia with chronic gastritis. Therefore, the associated myofibroblasts may already be exhibiting gene expression alterations.

Dysregulation of components of the Wnt signaling pathway is well characterized as early events in transformation and tumorigenesis, and therefore it would not be unexpected for this pathway to be already disrupted in ATM myofibroblasts. In support of this explanation, WNT5A upregulation was not identified in gCAM proteomes by Holmberg *et al*. (Holmberg et al., 2012), but transcriptional upregulation of WNT5A mRNA in gCAM compared to gNM was observed in a subsequent paper from the same group and shown to be functionally relevant (Wang et al., 2016). In further support of our observation of upregulated ROR2 in gCAM/pCAF, bone marrow-derived mesenchymal stem cells were shown to have significantly higher ROR2 mRNA than the gastric cancer cell line MKN45 (Takiguchi et al., 2016). Moreover, high ROR2 expression was identified in the stroma of 43% of pancreatic ductal adenocarcinoma (PDAC) tissues. However, this was not significantly different from matched tissues or benign pancreatic lesions, and the stromal cell type was not characterized (Huang et al., 2015). In addition, high stromal ROR2 expression was significantly associated with lymph node invasion and tumor stage in this study.

One of the most interesting observations from the results presented here is that a ligand WNT5A, together with the associated co-receptor ROR2, was transported from a signal-producing cell to a signal-receiving cell. This is the first time that direct transfer of a Wnt receptor has been reported, and it provides a paradigm shift in our understanding of how cells can respond to paracrine Wnt signaling. Fluorescently labeled ROR2/WNT5A complexes from producing cells were observed co-localizing in receiving AGS cells over considerable time periods and in most directly contacted cells. The receptor-ligand complexes were transferred via cytonemes in association with producing cell membranes, leading us to suggest that the tip of the cytoneme buds off following contact with a receiving cell and is endocytosed. In support of this hypothesis, we observed a similar phenomenon occurring in zebrafish embryos during development. In a parallel study by our group, quantitative fluorescent imaging via FCCS and FLIM-FRET was developed to provide evidence that zebrafish Ror2, and the relevant ligand Wnt5b, are significantly correlated in the producing cell, on cytoneme tips, and also in the receiving cell (Zhang et al., 2022). Moreover, cytoneme-dependent spreading of active Wnt5b/Ror2 complexes is critical for convergence and extension in the zebrafish gastrula. Taken together, we suggest a conserved and essential role for this signaling mechanism in both vertebrate embryogenesis and disease.

Despite being the first observation of Wnt receptor transfer to a signal receiving cell in a complex with its ligand, there is some precedent for this finding. Evidence suggests that exovesicles have a role in receptor transfer, including transport of Fzd10 mRNA in gastric and colorectal cancer (Scavo et al., 2019) and intercellular transfer of the oncogenic receptor EGFRvIII by microvesicles derived from glioma cells has been reported (Al-Nedawi et al., 2008). Although we do not rule out the possibility of exosome-mediated intercellular transfer of ROR2 in the findings presented here, the observation that paracrine JNK signaling correlates with the relative amount of producing-cell derived ROR2 in the receiving AGS cell provides evidence to indicate a more direct role for cell-to-cell transfer of activated ROR2 complexes, as all reporting cells would be exposed to soluble, secreted factors in this experiment. We clearly observed WNT5A/ROR2 complexes being transferred from cytoneme tips to receiving cells in both our parallel zebrafish model and the gastric cancer model presented here. Although this is the first time this transport mechanism has been characterized for ROR2, other receptors have been transported along cytonemes. Thickveins (the receptor for DPP) and EGFR have been observed moving in puncta along cytonemes in *Drosophila* (Hsiung et al., 2005, Roy et al., 2014). Moreover, confocal fluorescent microscopy of GFP-tagged FZD7 shows retrograde transport along cytonemes during chicken embryonic somite development (Sagar et al., 2015). FZD7 is trafficked along cytonemes that project towards the ligand-producing cells and then act as a transporter to bring ligands back to the cell. We have not observed a similar retrograde transport of WNT5A in association with ROR2 on producing cell cytonemes, but this possibility is not excluded. We have previously shown that ROR2 also co-localizes with other ligands and components of the Wnt signaling pathway on cytonemes (Mattes et al., 2018, Brunt et al., 2021, Routledge et al., 2022) including the canonical ligand Wnt8a. This raises the intriguing question of how the trafficking of different proteins to the cytoneme tip for transport is regulated and what molecular mechanisms determine which filopodia become Wnt signaling cytonemes. Moreover, the cytoneme contacts we have observed between gastric cancer cells appear to be less dynamic and more long-lasting than those in zebrafish development, leading to a more general question regarding how cytoneme target cells are determined.

In addition to observing the intercellular transport of ROR2/WNT5A complexes, our studies also reveal that pCAF2-derived, paracrine transport of ROR2 can induce JNK signaling in AGS receiving cells at a level that directly correlates with the amount of functional ROR2 transported. We also observed polarization of actin in receiving AGS cells towards ROR2 bearing cytonemes extended by pCAF2 cells as further evidence of induction of the WNT/PCP pathway. Despite an apparent decrease in overall AGS migration observed in a standard 2D wound healing assay with producing cells overexpressing ROR2 and WNT5A, we could attribute this to a directional migration cue that works to attract AGS cells towards cells expressing high levels of ROR2 using both an *in vitro* 3D invasion assay and an *in vitro* zebrafish model.

Previous studies have shown a role for MSC-derived ROR2 and WNT5A in autocrine induction of CXCL16, which in turn promotes the proliferation of MKN45 gastric cancer cells via CXCR6 (Takiguchi et al., 2016). In addition, the knockdown of CXCL16 in MSC suppressed the migration of MKN45 cells (Ikeda et al., 2020). However, bone marrow-derived MSC cells are a heterogenous pool of progenitor cells, and here we report specifically on the function of ROR2 in cancer-associated fibroblasts. Moreover, knockdown of ROR2 in MSC had a greater inhibitory effect on MKN45 migration than knockdown of CXCL16, suggesting a role for ROR2 supplementary to it’s role in CXCL16 upregulation. Furthermore, when AGS cells transfected with ROR2 and WNT5A are used as the signal-producing cells in our study, we report a similar level of paracrine JNK activation in the receiving AGS cells as when pCAF2 cells are the signal producing cells. In both, this experiment and in our parallel zebrafish model, it is unlikely that the same panel of chemokines and other soluble signals would be present. There are no obvious orthologues of CXCL16 or CXCR6 in zebrafish (Bussmann and Raz, 2015), although CXCL chemokines have been shown to have a role in migration at later stages of zebrafish embryonic development. In addition, blocking WNT5A-mediated ROR2 internalization suppressed gastric cancer tumor cell invasion and metastasis in WNT5A high but not WNT5A low cells (Hanaki et al., 2012), suggesting a direct role for ROR2 in activating the Wnt-PCP pathway rather than a chemokine signaling pathway. It has also been shown that filopodia formation mediated by ROR2 is required for WNT5A-induced cell migration in human melanoma cell lines (Nishita et al., 2006) further supporting our observation of a role for direct transport of ROR2 in inducing migration in AGS cells.

Finally, in addition to the novel intercellular signaling mechanisms identified here, we have also developed two novel assays for the study of cell migration. Firstly, bespoke PDMS microchambers were designed specifically to define, image, and characterize the molecular interactions between signal-producing and signal receiving cell populations. These represent an easily adaptable tool that allows closer and more reproducible contacts than traditional 2D scratch assays or wound healing chambers. Secondly, we developed and optimized an invasion assay in which AGS cells migrate through a transwell and into a 3D culture system containing fibroblast populations. This recapitulates the stroma surrounding gastric cancer cells, and by using a non-animal, non-carcinogenic derived hydrogel, the system is consistent between batches and provides the sensitivity required to dissect the contributions of different signaling components in the system.

In conclusion, we show that the WNT/PCP co-receptor ROR2 can be directly transported from cancer-associated fibroblasts to AGS cells, thereby inducing JNK signaling, actin polarization, and directional migration in the receiving cell in a ROR2 dependent manner. These observations were confirmed in an *in vivo* zebrafish model of cell migration reported in this paper and further supported by a parallel developmental zebrafish study that validates this novel WNT/PCP signaling mechanism. Taken together, these results increase our understanding of how gastric cancer cells can respond to their microenvironment and the molecular mechanisms that promote migration and metastasis.

## Acknowledgments

Research in the Scholpp lab, including support for S.R, is funded by the Medical Research Council (MRC)/UKRI through a research grant (MR/S007970/1), the BBSRC (Research Grant, BB/S016295/1 and an Equipment grant, BB/R013764/1), all awarded to S.S.. Further support is provided by the Living Systems Institute, University of Exeter, and C.Z. is supported by a Chinese Scholarship Council (CSC) studentship. This work was also generously supported by the Wellcome Trust Institutional Strategic Support Award (204909/Z/16/Z). Technical support from Agnieszka Kaczmar, Corin Lidle (Bioimaging facility), Francesca Carlisle (LSI tissue culture facility) and Raif Yuecel (Cytomics facility) is much appreciated.

## Author contributions

S.R. and S.S. designed experiments and wrote the manuscript. S.R. performed experiments and data analysis. C.Z. performed zebrafish experiments and analysis. M.F. provided the pCAF2 cell line used in this study. V.A. and F.G. designed and produced the microfluidics chips.

## Supplementary Figures

**Supplementary Figure 1.**
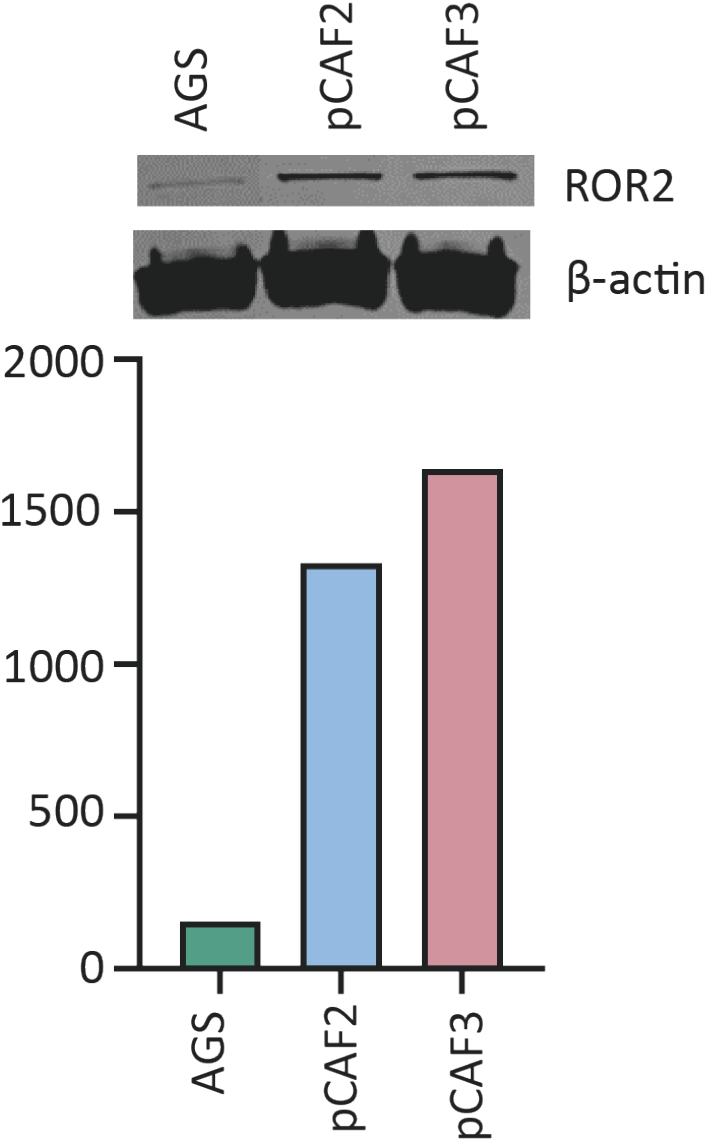
Correlation between ROR2 mRNA expression and protein expression was confirmed by Western blotting. The top panel shows ROR2 and β-actin bands in the indicated cell lines. The bottom panel indicates the intensity of the ROR2 band normalized to the intensity of the β-actin band.

**Supplementary Figure 2:**
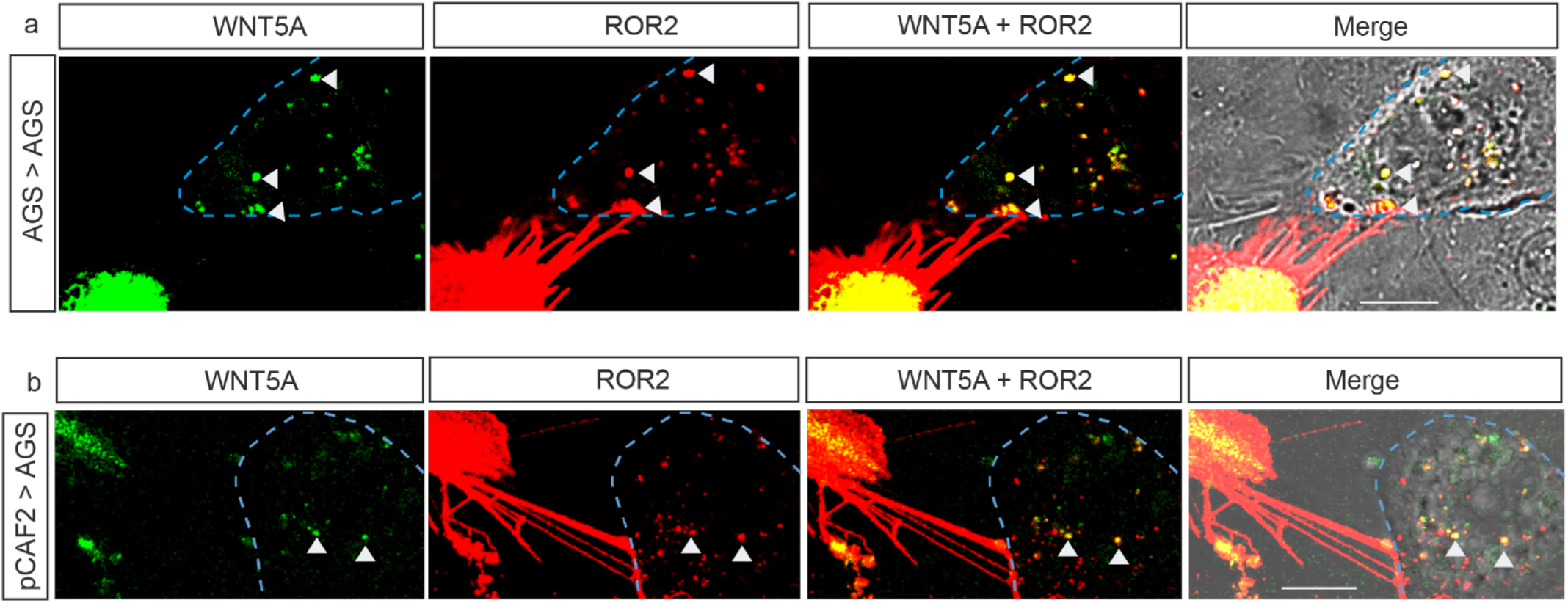
ROR2/WNT5A complexes are transferred from producing cells to receiving cells via cytonemes. **a**. WNT5A-GFP and ROR2-mCherry were overexpressed in producing AGS cells, and co-cultured with untransfected receiving AGS cells for 24 hours prior to imaging with confocal microscopy. Blue dashed line indicates the position of the receiving cell as detected by the brightfield image. Co-localising complexes in the receving cell are indicated with a white triangle. **b**. WNT5A-GFP and ROR2-mCherry were overexpressed in producing pCAF2 cells, and co-cultured with untransfected receiving AGS cells for 24 hours prior to imaging with confocal microscopy. Blue dashed line indictaes the position of the receiving cell as detected by the brightfield image. Scale bar indicates 10μm.

**Supplementary Figure 3:**
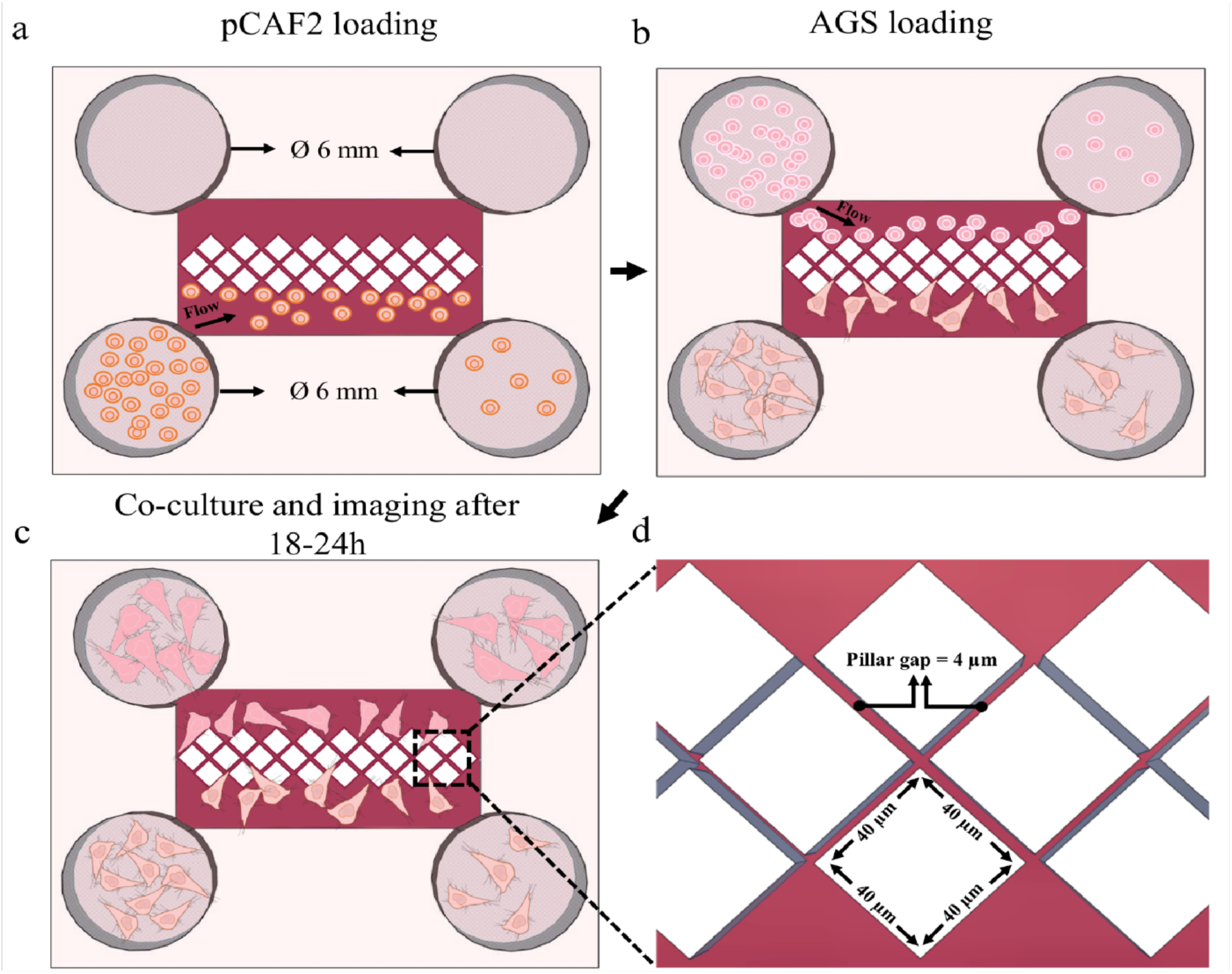
Schematic of the microfluidic device to study pairwise cytonemal communication. **a**. On one side, pCAF2 cells are pipetted into large 6 mm wells from where they flow towards the pillar array. **b**. After 24h incubation, the AGS cells are introduced into the other side, allowing them to be placed at close distance to the pCAF2 cells. **c**. After co-culture for 18-24 hours, cells are imaged at high-resolution. **d**. A planar microfluidic chamber is divided into two sides of equal volume by a linear array of diamond shaped posts. The 4 µm pillar gaps are designed such that pCAF2 cells cannot migrate through the gaps whereas filopodia can.

**Supplementary Figure 4:**
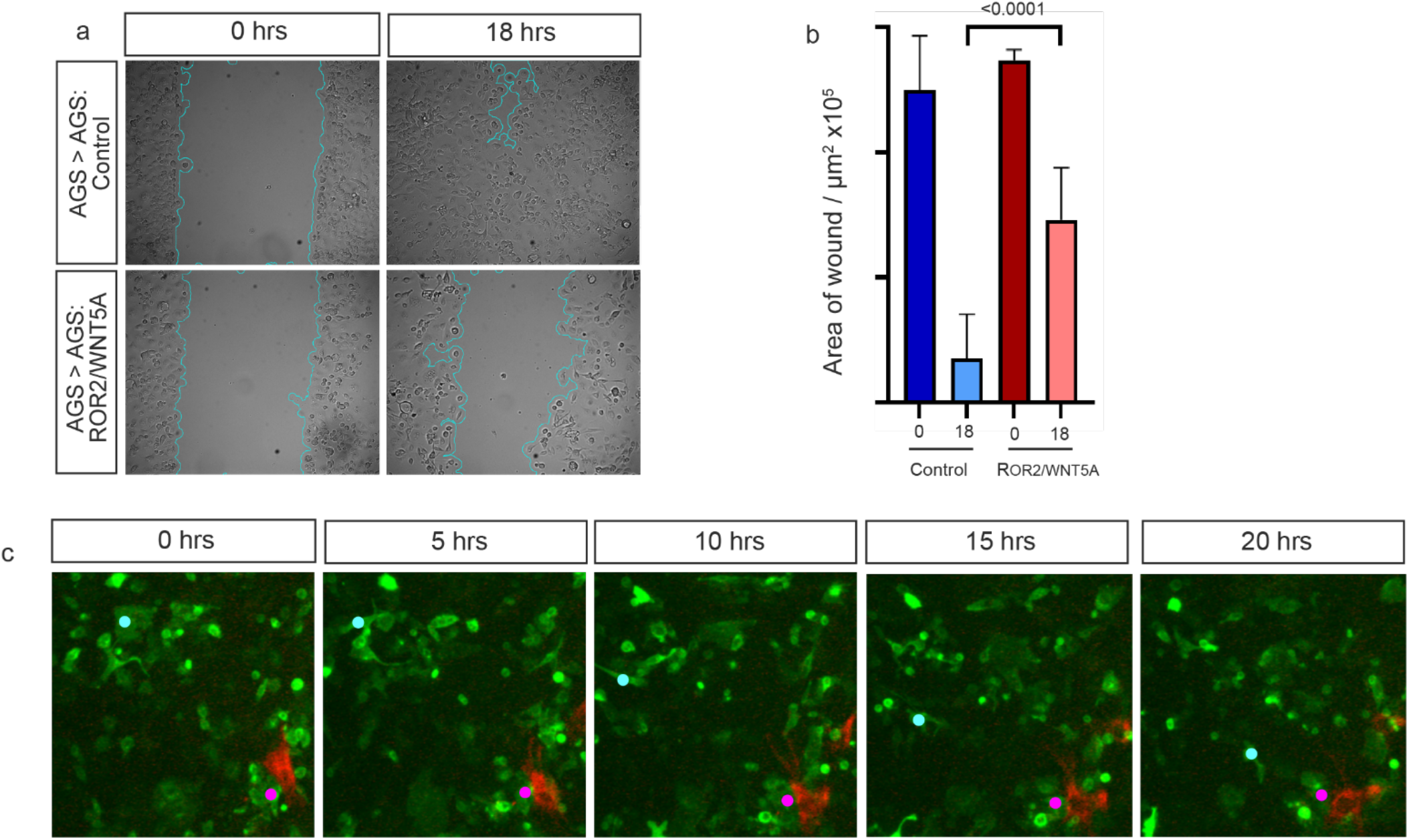
ROR2/WNT5A overexpression induces a directional migration in AGS cells. **a**. Control AGS or AGS cells transfected with Ror2 and Wnt5a expression plasmids were cultured in Ibidi wound healing chambers for 24 hours prior to mitomycin C treatment and release. Dishes were imaged at release (0 hrs) and 18 hrs later. **b**. Area of the wound at 18 hours was measured at the start and end of the experiment. **c**. Time lapses images taken at the indicated times of pCAF2 expressing ROR2 mCherry (red cells) co-cultured with AGS cells expressing membrane GFP. An individual AGS receiving cell in close proximity to a pCAF2 cell is highlighted with a pink dot, whereas an individual AGS receiving cell in that starts at a distance to pCAF2 cells is highlighted with a light blue dot

